# Innovative Research Experiences for Underrepresented Undergraduates: A Collaborative STEM Research Program as a Pathway to Graduate School

**DOI:** 10.1101/2023.01.03.522432

**Authors:** Gokhan Hacisalihoglu

**Affiliations:** Biological Sciences Department, Florida A&M University, Tallahassee, FLORIDA 32307, USA

**Keywords:** Biology, Plant science, Undergraduate students, STEM, Mentoring, Gateway

## Abstract

This paper aims to describe, reflect on, and explore the perceptions of underrepresented undergraduate researcher students towards the plantREU2 internship program of two universities (an HBCU and an R1 university) in Florida, USA. The plantREU2 internship program resulted from a collaboration between UF and FAMU and was located in UF main campus in Gainesville, Florida. A total of 17 students completed 10-week summer projects in plant biology. The program (PlantREU2) had a strong record of success. Over 40% of plantREU2 students co-authored a journal publication and received travel awards to attend Maize Genetics Conference. Furthermore, plantREU2 participants were significantly graduated within six years. The underrepresentation of minorities in STEM is a critical challenge. The findings of this study can be adapted similarly for underrepresented undergraduates. The vast majority of interns enrolled in post-graduate programs could therefore be a model to engage traditionally underrepresented students in STEM disciplines.

## INTRODUCTION

There are multiple positive benefits from engaging students in undergraduate research experiences (UREs). Undergraduate research increases learning, research skills, and retention in a large diversity of academic fields (McDevitt et al., 2016). In STEM disciplines, there are several models to engage students in research, including course-based and one-on-one mentoring experiences in active research laboratories. In one-on-one UREs, students typically work with committed mentor scientists to improve their research skills and knowledge in a specific scientific discipline (Hacisalihoglu et al., 2022; 2021; 2020; 2018; 2016; 2010; 2007). Some programs seek to engage students in long-term research mentoring from lower division students, while others provide short-term UREs.

Undergraduate research is a significant factor in promoting graduation with a science degree, enrollment in STEM post-graduate study, and working in science-based fields (Hernandez et al, 2018). There is evidence that long-term UREs provide greater benefits to students regarding outcomes in science education and participation in the science workforce (Hernandez et al., 2018). However, long-term UREs may not be practical for underrepresented minorities when resources and research efforts are limited at smaller institutions.

Increasing the participation of minorities is a major goal of federal research funding agencies to develop a diverse STEM workforce. At historically black colleges and universities (HBCUs), comprehensive STEM interventions include myriad strategies to engage and promote student persistence towards post-graduate study. For example, the Morehouse College Hopps Scholars Program significantly increases the enrollment of African American male students in advanced degree STEM education. However, these programs require inter-institutional UREs for students to become well-prepared for graduate school. Inter-institutional URE programs face challenges in engaging mentors and students to train away from their home institutions (Morales et al., 2017)

The objective of this program is to create a summer research experience program between the University of Florida (UF) and Florida A&M University (FAMU) for underrepresented undergraduates to support their research training in plant genomics and their overall next-level career development.

## MATERIALS AND METHODS

### PlantREU2 Program

Improving crop plants and their seeds through novel analysis techniques and plant stress tolerance is continuing to be an important task in agriculture. The plantREU2 internship program resulted from a collaboration between UF and FAMU and was located in UF main campus in Gainesville, Florida. FAMU is a historically black college or university (HBCU) within the state university system together with UF. True to its motto of Excellence with Caring, FAMU is located in Tallahassee, Florida and committed to exemplary teaching of underrepresented minorities. The plantREU2 program started in the Summer 2007, and FAMU undergraduate students were recruited through visiting lectures of UF faculty at FAMU campus during the Spring semesters based on student GPA, plant biology interest, and letters of recommendation. A total of 17 undergraduate students from FAMU biology pre-professional majors participated in plantREU2 program between 2007 and 2018. The internship duration was ten weeks during the summer months, and each student received a bi-weekly stipend. The plantREU2 scholars first went through lab research training and lab safety, followed by directed research under a mentor on various research projects. The students gave an oral presentation of their findings on the last day of their internship. In addition to research work in the lab and field, plantREU2 scholars attended plant science seminars, annual maize genetics conference, as well as analyzed their own data.

### Examples of plantREU2 Projects

Our plantREU2 program involved undergraduate students in helping them learn plant genomics by involving in laboratory and field activities. The followings are summaries of the NSF-funded scholar projects examples developed under the plantREU2 summer program between FAMU and UF (Fig. 1).

**Figure 1.**
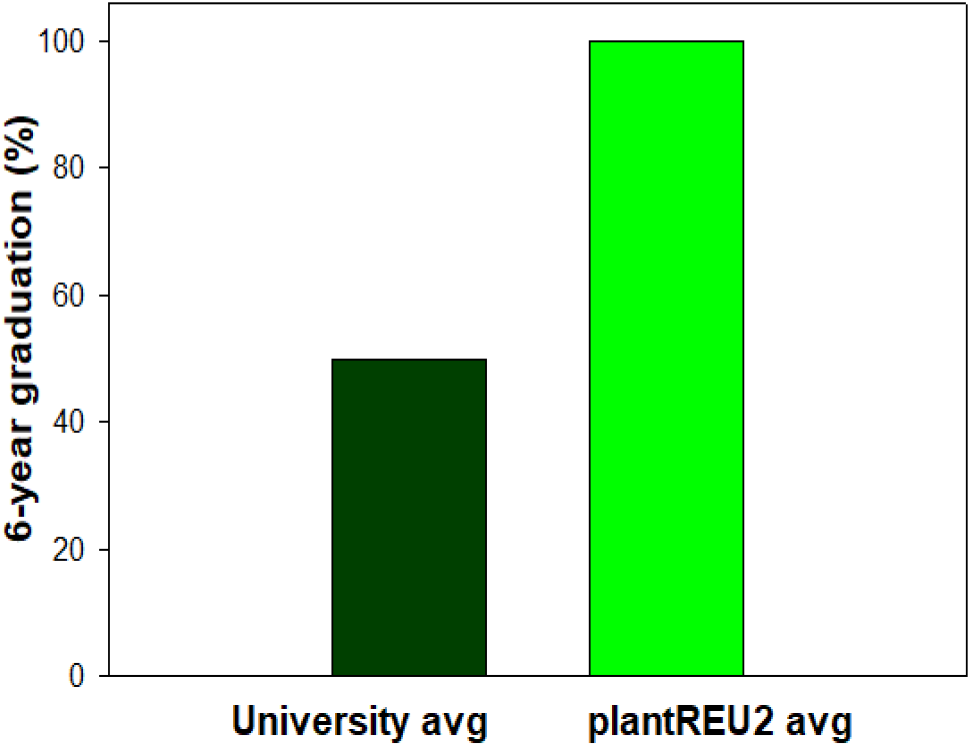
PlantREU2 students experiencing plant science research during the internships. Six-year graduation rate averages for the university and plantREU2 scholars.

#### Example Project 1

Predicting seed composition with single kernel NIRS. Organic compounds naturally absorb near infrared (NIR) light due to their functional groups. Consequently, NIR allows chemical inferences non-invasively by sample penetration. In this project, NIR spectra were collected from maize, bean, or soybean seeds using an automated single kernel analyzer. The seed samples were then analyzed to determine their weight, oil, protein, starch, and density levels. The results of this project showed the potential of chemometric and non-invasive prediction of seed composition in maize (Spielbauer et al., 2009), common beans (Hacisalihoglu et al., 2010), soybeans (Hacisalihoglu et al., 2016), and peas (Hacisalihoglu et al., 2020).

#### Example Project 2

How to cultivate maize in cold weather: Warm temperature is essential for plant growth and development. Cold stress results in reduced emergence and productivity in crop plants. In this project, machine learning and video-based phenotyping were utilized to study genetic variation in maize NAM parental lines for cold tolerance. Results showed broad genetic variation for tolerance to cold stress (-10 C) was observed in maize (Hacisalihoglu et al., 2018).

#### Example Project 3

Analysis of NIR dosage effect mutants in maize: This research project aimed to explore the heritability of dosage effect mutants in maize. A single kernel instrument was used to screen for weight and seed quality mutants. The main focus of the project was the small number of mutants to identify loci that show heterozygous changes such as 2:1 or 1:1:1 ratios of differential weight or quality classes. The results suggested that dek mutants cause dosage effect irrespective of the parent of origin.

#### Example Project 4

Development of micro-CT x-ray tomography to analyze density and shape: Seed size and density traits play an important role in crop yield. We explored the potential of micro computed X-ray tomography (micro-CT) as a rapid way to accurately measure these traits. Results of this project showed that micro-CT is a powerful technique for analyzing especially density, volume, area, length, and width of individual maize (Gustin et al., 2013) and soybean seeds (Hacisalihoglu et al., 2016, Hacisalihoglu, 2022; 2021; 2020; 2011; 2007; Hacisalihoglu and Armstrong, 2022; Hacisalihoglu and Strickland, 2019; Hacisalihoglu and Settles, 2017; Hacisalihoglu and Settles, 2013; Hacisalihoglu and Ross, 2010; Hacisalihoglu and Vallejos, 2005).

## RESULTS

### Outcomes and student accomplishments

In general, plantREU2 summer internship program has led to oral and poster presentations, co-authored scientific publications, awards, as well as enrollment in graduate or professional schools. Since 2007, seventeen underrepresented undergraduate students have participated in the plantREU2 internship program. A total of 17 (100%) participating student scholars gave scientific presentations. Seven of the scholars co-authored peer-reviewed publications (a total of 41%) (Spielbauer et al., 2009; Hacisalihoglu et al., 2010; Gustin et al., 2013; Hacisalihoglu et al., 2016, 2018, 2020). Five of the scholars received very competitive travel awards to attend three maize genetic conferences. Several plantREU2 scholars have annually contributed to the FAMU student research forum and won the best poster prizes. Furthermore, nine (82%) plantREU2 scholars have successfully enrolled in graduate or professional schools. The remaining two students are in the process of preparing for graduate schools (Table 1).

**Table 1.** Academic outcomes of summer research program between UF and FAMU.

|  |  |
| --- | --- |
| Number of student scholars enrolled in the summer research internship | 17 |
| Number of student scholars gave presentation | 17 |
| Number of student scholars co-authored a scientific publication | 7 |
| Number of student scholars received conference awards | 5 |
| Number of student scholars who enrolled graduate or professional schools | 14 |

We explored whether 6-year graduation rates differed between plantREU2 students and non-plantREU2 students. Our results showed that plantREU2 internship students exhibited significantly (2-fold) higher 6-year graduation percentage compared to university 6-year graduation percentage (Fig. 1)

We developed a voluntary survey to evaluate the experiences and current career status of the plantREU2 scholars between 2007 and 2019. In addition to academic outcomes, the success of the plantREU2 program is best conveyed by written comments as summarized in Table 2.

**Table 2.**
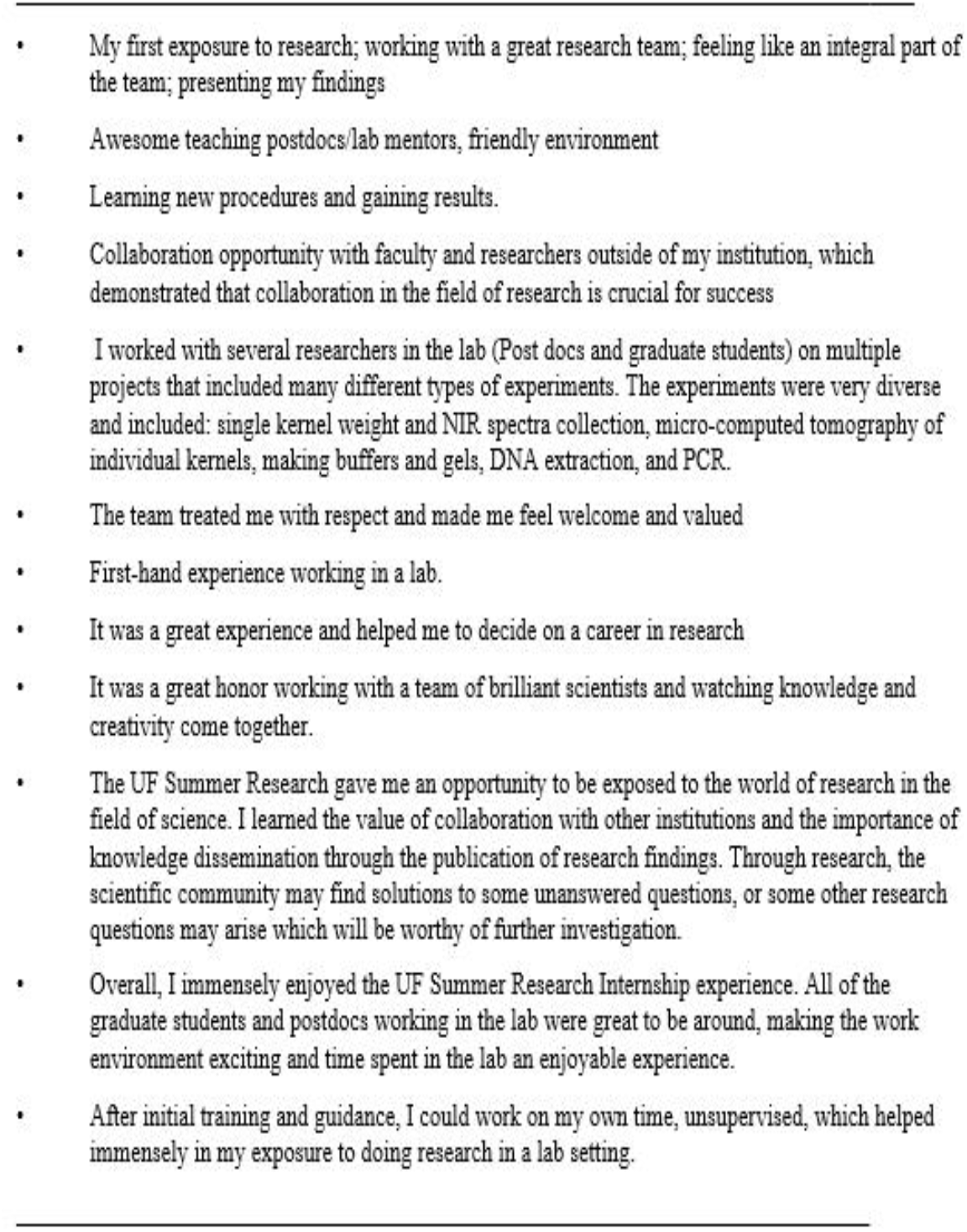
Evaluation and sample responses and written comments from plantREU2 scholars.

## DISCUSSION

We believe that the plantREU2 program described here is an innovative and successful research training program and introduced tra-ditionally underrepresented biology students to research and critical thinking skills necessary for the workplace of the 21st century.

The plantREU2 scholars spent an average of 10 weeks working as full-time research interns on projects ranging from high throughput phenotyping to cold tolerance and dosage effect mutants in maize. At the end of their sum-mer internship, plantREU2 scholars shared their research findings via exit seminars, con-ference presentations, and scientific publica-tions. Furthermore, most of plantREU2 scholars end up enrolling in graduate or professional schools afterwards.

In conclusion, diversifying the plant biology workforce will keep the United States competi-tive in the world economy, and this is one of the most unique summer undergraduate research programs for HBCU students. Finally, the plantREU2 program could be successfully im-plemented to create a gateway to STEM and plant science careers. Finally, this article introduces an effort to relate a case study of a successful partnership between an HBCU and an R1 research institution.

## ACKNOWLEDGEMENTS

We gratefully acknowledge National Science Foundation (USA) for funding. We would like to acknowledge the students for being part of this REU program, Dr. A. M. Settles for exceptional collaboration, host, and making this REU program happen, and his many years of leadership and advocacy for this partnership, as well as anonymous reviewers and the editor in providing detailed feedback.

## Notes

### Competing Interest Statement

The authors have declared no competing interest.

